# A stable, high refractive index, switching buffer for super-resolution imaging

**DOI:** 10.1101/465492

**Authors:** Tobias M.P. Hartwich, Kenny Kwok Hin Chung, Lena Schroeder, Joerg Bewersdorf, Christian Soeller, David Baddeley

## Abstract

dSTORM super-resolution imaging relies on switching buffers to enable dye molecules to enter and exit a metastable dark state. Current buffers have a very limited shelf life of approximately 1 day and poorly match sample refractive index, impacting negatively on measurement reproducibility and image fidelity. We present a buffer based on chemical, rather than enzymatic, oxygen scavenging which exhibits dramatically improved stability, switching speed, contrast, and index matching.

---

Super-resolution techniques such as FPALM, PALM, and STORM^1-3^ circumvent the optical resolution by mapping out the positions of individual fluorescent molecules attached to a structure of interest. This is achieved by ensuring that only a sparse subset of molecules are visible in any given frame, such that individual molecules are well separated and can be clearly distinguished. Under these conditions, the position of each imaged molecule can be determined with a precision that depends principally on the number of photons collected from that molecule^4^ rather than the wavelength of light used. By imaging many different subsets of molecules over tens of thousands of image frames, a resolution beyond the diffraction limit can be achieved^5^.

The ability to image sparse subsets of molecules requires the majority of molecules to be in a transient non-fluorescent (dark) state at any given point in time. One of the most common ways to achieve this is by pumping the fluorophores into a long-lived, dark state using intense laser excitation and a reducing imaging-buffer that contains a primary thiol such as 2-mercaptoethanol (2-ME) or cysteamine (MEA). This technique, commonly called dSTORM^6^,but also known as e.g. GSDIM^7^ or RPM^8^, relies on a special imaging buffer both to reduce the amount of permanent photobleaching and to facilitate the reduction of fluorophores into the dark state. The most commonly used switching buffer, consisting of glucose oxidase, catalase, glucose and MEA in PBS or Tris buffer^9, 10^, has severe drawbacks.

One important limitation of current buffer systems is that they are typically aqueous, with a refractive index of ~1.33 and not index matched to either the high NA immersion objectives (~1.51) commonly used for superresolution imaging or to the refractive index of intracellular compartments (~1.38-1.45)^11^. A slightly modified buffer system that consists of 80-90% glycerol and MEA^12^ enables better index matching and is thus preferable for imaging thicker samples. Unfortunately, the enzymatic activity of glucose oxidase is severely impaired in glycerol^13^, resulting in little to no active oxygen scavenging. Surprisingly, the oxygen content of glycerol-containing thiol buffers is low, even when enzymes are omitted, implying that the oxygen scavenging in high index buffers occurs largely due to consumption of the thiol itself^14^,assisted by a lower oxygen mobility^8^ resulting from the increased buffer viscosity.

The most critical limitation of current buffers is however a lack of stability, with commonly used buffers showing a rapid and unpredictable loss of function over the course of a few hours - even when stored at 4° C. Multiple factors contribute to this loss of function including the fact that the oxygen scavenging reaction of the glucose oxidase-catalase system acidifies the buffer over time^15, 16^, and that the thiol is readily oxidized and consumed by dissolved oxygen^14^. Importantly, the oxidation of thiol components still occurs at oxygen concentrations well below the lower limit of enzymatic scavenging systems. In addition to the inconvenience of having to make fresh buffer before each imaging session, the unpredictability of the decay introduces an uncontrolled parameter into all measurements, complicating efforts to reproduce and quantify imaging results.

Here we demonstrate an alternative imaging buffer for dSTORM consisting of 80-90% glycerol, MEA, and sodium sulfite (henceforth referred to as the ‘sulfite buffer’). Sodium sulfite is a well-established chemical oxygen scavenger^17^ that is the main ingredient of ‘zero-oxygen’ calibration solutions^18^, and widely used as a food preservative^19^. It works well in a high glycerol content buffer and avoids acidification of the imaging buffer due to its simple reaction stoichiometry: 2*Na_2_SO_3_* + *O_2_* -> *2Na_2_SO_4_*. Note that although cobalt ions are often used as a catalyst, we found the scavenging reaction proceeded rapidly even in the absence of Co^2+^. Sulfite based scavenging reduces the dissolved oxygen levels down to the limit of our oxygen meters detection sensitivity, in contrast to glucose oxidase which plateaus at ~.9%^20^.

One of the shortcomings of the conventional buffers mentioned above is the lack of long term stability. Using a buffer that is only a few days old, even when kept at 4°C, leads to dramatically inferior results. In contrast, we found that we could keep the ‘sulfite buffer’ at room temperature for several weeks with no discernible difference in the quality of the reconstructed images. This is illustrated in Figure 1 b & c, which show images of Alexa 647 labeled α-tubulin in Cos7 cells recorded with fresh and 28 days old buffer respectively. A quantification of the number of blinking events observed per area of microtubule labeling (Fig. 1a, see supplementary materials for details) confirms the qualitative results. An imaging buffer containing only 90% glycerol and sodium sulfite without the addition of a primary thiol was not suitable for dSTORM imaging, indicating that the sulfite acts as an oxygen scavenger and preservative rather than as a reducing agent itself. This preservative action is likely due to the extremely low oxygen levels, which will effectively inhibit the spontaneous oxidation of thiol, but may also in part be due to the fact that the pH of the buffer remains stable during scavenging.

**Figure 1.**
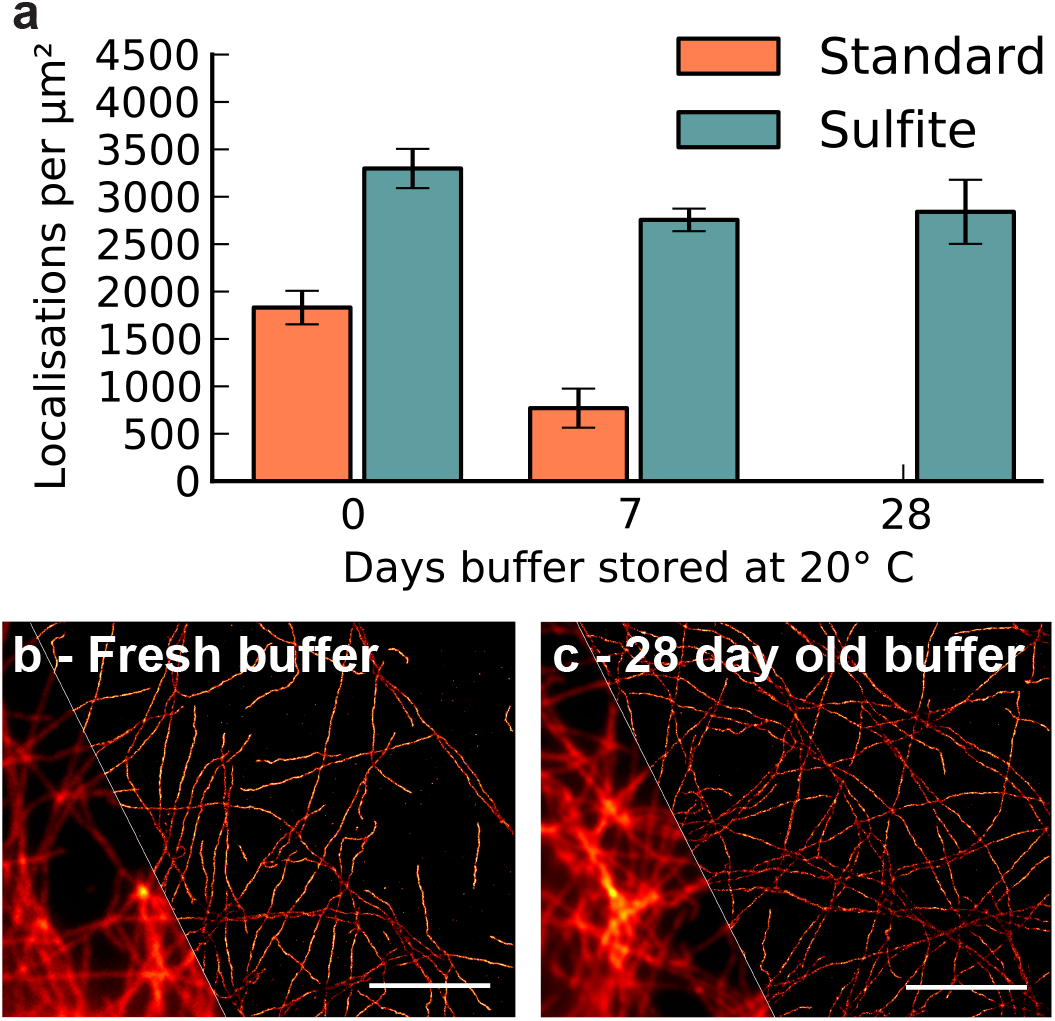
Effect of buffer age on superresolution imaging. (a) The localization density falls quickly off as standard buffers age, but is maintained when sodium sulfite is included. This results in a preservation of imaging performance (b, c), as seen in dSTORM images of α-tubulin labeled with Alexa 647 using fresh and 28 days old sulfite containing buffer. Diagonal insets show the diffraction limited images, and the scale bars are 2 µm.

Figure 2a shows kymographs of two dSTORM data sets imaging different cells in the same sample using the ‘MEA-glycerol buffer’ (top) and the ‘sulfite buffer’ (bottom). All other conditions are the same. From the kymographs we can see that molecules in the ‘sulfite buffer’ exhibit much shorter fluorescence bursts, or ‘blinks’ before transitioning back into the dark state than those imaged using the ‘MEA-glycerol buffer’, and that the background in the ‘sulfite buffer’ is significantly lower. This is quantified in Figure 2b, which shows that the on-state life time of Alexa 647 in the ‘sulfite buffer’ is negatively correlated to the sulfite concentration. We attribute the reduction in blink time to an increased effective thiol concentration as well as a removal of the triplet-quenching effect of dissolved oxygen. The slight increase of the life time between the ‘MEA-glycerol buffer’ (0 mM sulfite in Fig. 2b) and the ‘sulfite buffer’ with 1 mM sulfite can be tentatively explained by the 1 mM concentration being enough to preserve thiol concentrations, but not yet enough to significantly reduce the effect of oxygen mediated triplet quenching. The reduced background results from an increase in the proportion of molecules in the dark state at any given time, leading to less out of focus fluorescence and consequently a better signal to noise ratio for each event (Fig. 2a). As localization precision depends strongly on background levels, this reduction in background (Supplemental Fig. 1a) readily compensates for the slight decrease in the number of photons collected per molecule (Supplemental Fig. 1b) due to the shorter on-state life-time, such that the localization precision improves with increasing sulfite concentration (Fig. 2c). This effect is already apparent in the comparatively thin specimens imaged here and is expected to become more significant for thicker samples for which the initial background is already high. The observed dependence of the on-state life time on sulfite concentration offers an additional experimental ‘handle’ with which to fine tune blinking times to match camera frame rates and to allow an increase in acquisition speed.

**Figure 2.**
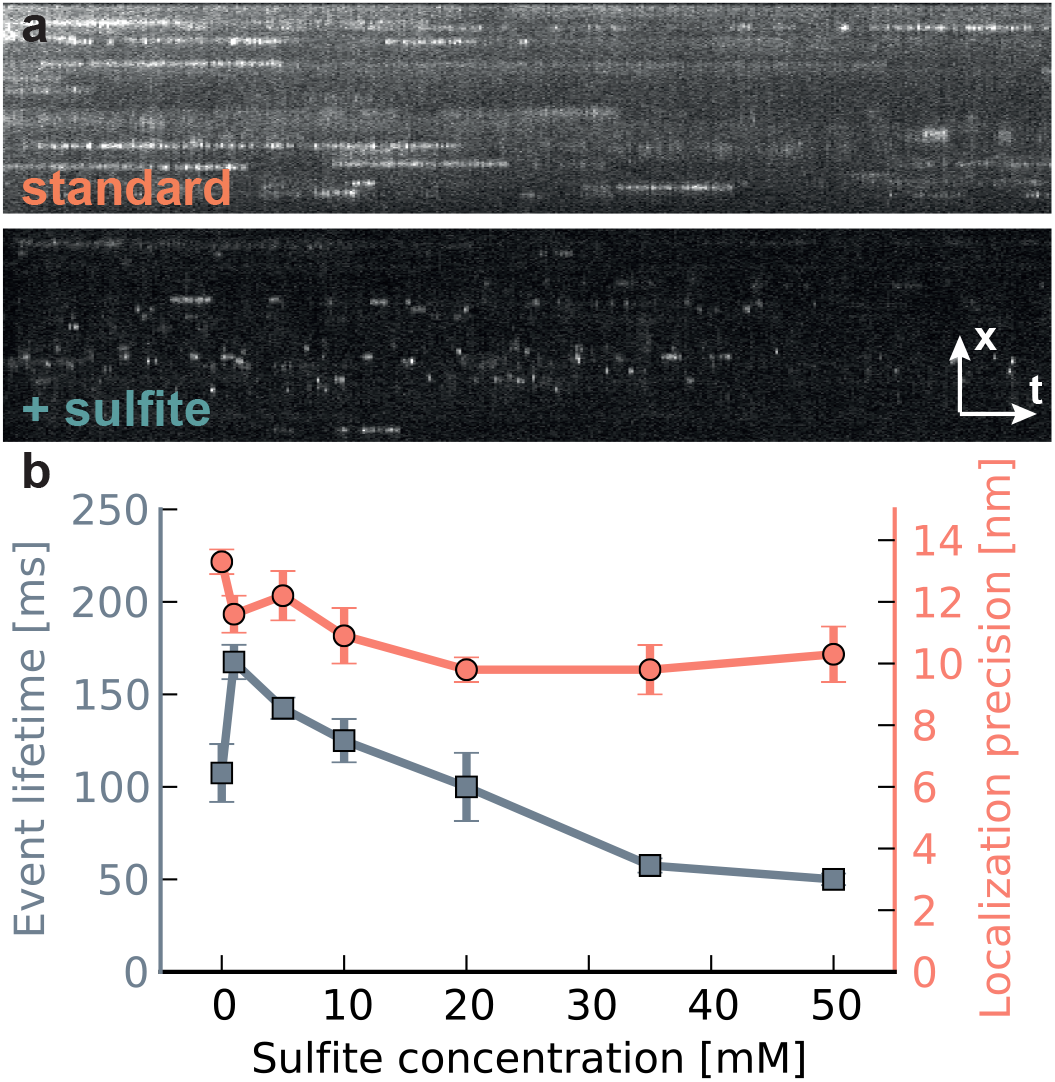
Effect of sodium sulfite on single molecule switching. Kymographs of dSTORM data sets (a) recorded without and with sulfite show that the addition of sulfite results in a reduction in fluorophore on-time and an improvement in contrast. Quantifying these results (b), shows that both the on-state lifetime and the localization error decrease with increasing sulfite concentration.

In this paper we have focused on the use of sulfite in a 90% glycerol buffer as there are, to our knowledge, no other viable oxygen scavengers for high index buffers. Our buffer has a refractive index of ~1.45, resulting in a 3-fold reduction in spherical aberration over aqueous buffers and significantly improving imaging of structures which are not in the immediate vicinity of the coverslip. Sodium sulfite can, however, also be used in aqueous buffer (Supplemental Fig. 2) with a similar reduction in background and improvement in localization precision. In addition to Alexa647, we have tried a number of other dyes, including Alexa 680, Alexa 700, Cy3B and Alexa 594 which all switch well in sulfite containing buffer.

In summary, we found that the chemical oxygen scavenger sodium sulfite improves fluorophore switching whilst also acting as a preservative for the MEA in the buffer, providing long-term stability. Its use instead of the enzymatic glucose oxidase-catalase system results in shorter fluorescence bursts, lower background, higher event densities, and better overall contrast without compromising on localization precision. We anticipate that the dramatically improved stability of the sulfite containing buffer will not only simplify the imaging process but more importantly enhance reproducibility between imaging sessions.

